# Genetic barcoding uncovers the clonal makeup of solid and liquid biopsies and their ability to capture intra-tumoral heterogeneity

**DOI:** 10.1101/2025.06.12.659267

**Authors:** Antonin Serrano, Tom S Weber, Jean Berthelet, Sarah Ftouni, Farrah El-Saafin, Verena C. Wimmer, Kelly L. Rogers, Elgene Lim, Emmanuelle Charaffe-Jauffret, Christophe Ginestier, David Williams, Frederic Hollande, Belinda Yeo, Sarah-Jane Dawson, Shalin H. Naik, Delphine Merino

**Author notes:** These authors equally contributed to this work (Antonin Serrano, Tom S Weber). Joint senior authors (Sarah-Jane Dawson, Shalin H. Naik, Delphine Merino).

## Abstract

Intra-tumoral heterogeneity (ITH) is fueling tumor progression in breast cancer, as specific clones present within a tumor may have a selective advantage to colonize distant organs and escape therapy. Accurate sampling of ITH is therefore a pressing challenge in clinical oncology, to adequately predict recurrence and inform rational and personalized therapies. Here we used cellular barcoding to track the spatio-temporal composition of human breast cancer clones in six preclinical models – across two cell lines and four patient derived xenografts (PDXs). This allowed direct side-by-side quantitative comparison not only of intra-tumor clonal composition, but also of how that composition was reflected in needle biopsies and cell-free DNA (cfDNA). These analyses highlighted several clinically relevant findings. First, the use of orthogonal genetic and optical barcoding revealed that clonal diversity in the center of non-necrotic primary tumors was significantly higher compared to their periphery. Second, cfDNA barcode analysis suggested that DNA ‘shedding’ in the vasculature varied largely, not only depending on necrosis and tumor burden, but also between models. Third, combining information captured in both solid and liquid biopsies can provide a more robust assessment of tumor clonal composition. Taken together, these results showcases the utility of these barcoded models to optimize the use of solid and liquid biopsies as surrogate markers of ITH.

## Introduction

Breast tumors are composed of a complex patchwork of cancer clones, each with differential abilities to invade locally and distally^1^. While the exact mechanisms underpinning metastasis progression and drug resistance are not yet fully elucidated, intra-tumoral heterogeneity (ITH), including genomic^2^ (DNA mutations) or non-genomic^3^ (e.g. epigenetic and transcriptomic) heterogeneity, correlates with disease progression^4,5^. However, the contribution of individual clones to disease progression is not necessarily correlated with their size in the primary tumor, as low-frequency clones can be highly metastatic in distant organs^6–9^. These observations highlight the need to comprehensively assess the clonal heterogeneity of primary tumors to better predict and treat recurrence.

Currently, breast cancer diagnosis and prognosis testing are based on solid biopsies or specimens collected during surgery. The status of ER, PR and overexpression of HER2, as well as markers of aggressiveness, are routinely contributing to identifying the subtype and grade of the disease, which in turn guides therapeutic decision^10^. However, the level of heterogeneity captured in biopsy is still unclear. Yet, since most efforts in precision oncology are focused on identifying biomarkers predictive of outcomes and drug response for individual patients^11,12^, methods of biopsy must provide a precise representation of the tumor complexity. Thus, a full appraisal of the different types of biopsies, and how accurately they reflect tumor heterogeneity and predict therapeutic outcomes is required in the design of personalized therapies.

Liquid biopsies from blood specimens are non-invasive and measure the DNA sequence of cell-free DNA (cfDNA) and circulating tumor DNA (ctDNA). This form of biopsy has emerged as a promising option to detect and monitor tumor progression and treatment responses in cancer patients^13–16^. Previous studies indicated that ctDNA is able to capture the genomic ITH of patient tumors^17^, whereas core needle biopsies or fine-needle aspiration may not be entirely representative of the genomic landscape of tumors^18–20^. Furthermore, ctDNA biopsies have the advantage of being less invasive and more likely to represent the overall heterogeneity of the disease when multiple lesions cannot be easily biopsied simultaneously in clinical settings^21^.

While differences between biopsy methods have previously been investigated^19,22,23^, most studies focus on a few driver mutations (TP53, MEK1, KRAS, EGFR) rather than characterizing overall clonal heterogeneity of the disease. Comparison of ctDNA with needle biopsies in gastrointestinal cancer indicated that ctDNA was able to capture resistance alterations not found in the matched tumor biopsies in 78% of cases^24^. This suggests that ctDNA might be a better surrogate of genomic heterogeneity compared to solid biopsies. However, the ability of cancer clones to shed ctDNA into the bloodstream might depend on the level of necrosis in tumors^25^ and on the intrinsic characteristics of the clones^26^. Furthermore, the recent development of single-cell RNA sequencing (scRNA-seq) analysis and drug prediction from transcriptional signatures, rather than mutational status, highlights the utility of capturing and analyzing intact cells in cancer research and precision medicine. This is particularly relevant for non-genetic cancer alterations. Therefore, improvements in the collection of fresh tissues from needle biopsies would be of clinical utility in diagnostics and personalized medicine.

Cellular barcoding strategies using genetic barcoding^9,27–29^ or optical barcoding^30,31^ have emerged as powerful tools to study the heterogeneity of breast cancer primary tumors and metastases. These lentiviral-based labeling strategies allow the labelling of individual cancer cells with unique genetic or optical tags, respectively^32–34^. As these tags are stably integrated into the genome of the transduced cells and transmitted to their progeny, clonal fate can be monitored in large populations of cells, regardless of the molecular profiles of the clones and their spatio-temporal evolution. These preclinical models enable a comprehensive analysis of the clonality of the disease in multiple mice, tissues, and conditions (i.e. in absence of treatment, at multiple times), and therefore provide quantitative and qualitative information that can’t be easily captured using patient samples. Here, we used cellular barcoding to label human breast cancer tumors and explore the clonal repertoire captured in solid and liquid biopsies, by identifying the barcodes present in ex-vivo needle biopsy samples and in the plasma of tumor-bearing mice. This analysis provided a unique opportunity to comprehensively assess the ability of different sampling methods to capture ITH.

## Results

### Clonal density is higher in the center of primary tumors

Previous studies using multi-region sequencing on patient samples have shown that clones grow as patches within primary tumors^18,20,35–38^. To better understand the spatial distribution of clones in primary tumors and how this might affect the quality of solid biopsies, a previously established simulation model was adapted^39^ **(Fig. 1a)**. This model predicts the 3D growth of clones based on three cellular parameters: birth rate, death rate and mobility. Mimicking previous cellular barcoding transplantation experiments^9^, cells were given an identity at the start of the simulation (in this case, using a virtual tag), thereafter inherited by their respective progeny. The simulation of clonal growth confirmed the clonal ‘patchiness’ of the tumors previously observed using genetic barcoding^9^. Interestingly, the modeled clonal density in the center of the tumor was higher compared to the periphery **(Fig 1b)**. Indeed, the virtual dissection of these tumors into five pieces indicated that the center of the tumor (piece E) exhibited a significantly increased number of clones. This heterogeneous topographical distribution in clonal density observed throughout the simulated tumors raised an interesting question. Namely, does this hold true in an in vivo situation, where additional factors like mechanical pressure and constraints imposed on a tumor growing in a mammary fat pad could influence clonal distribution?

**Figure 1.**
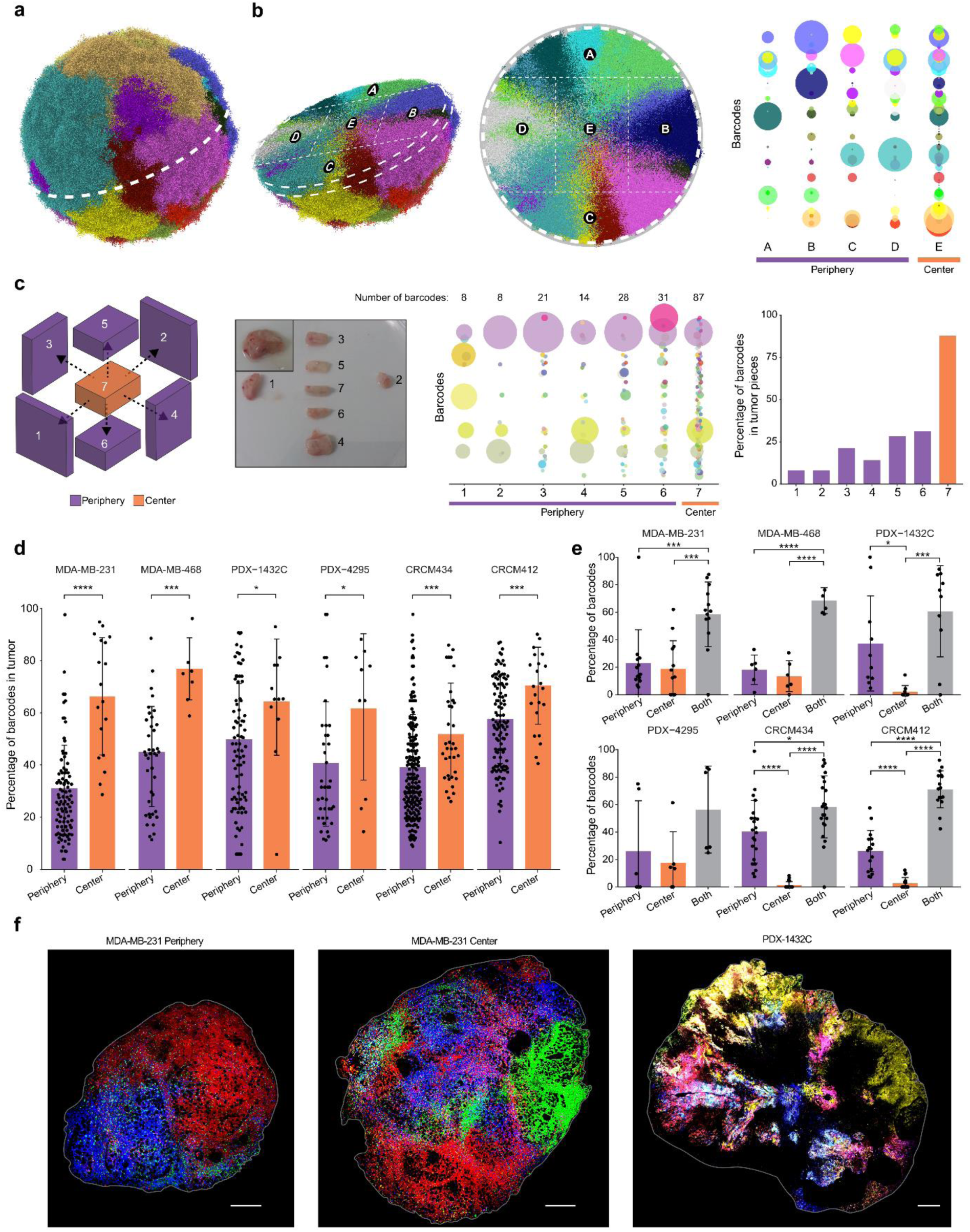
Spatial heterogeneity in primary tumors. **a)** 3D visualisation of clonal growth by simulation, based on virtually barcoded cancer cells. **b)** Dissection of the 3D virtual tumor into pieces (left), and representation of the barcode distribution in each piece (right). Each circle represents a barcoded clone. Its size correlates with the barcode frequency in each piece. **c)** Dissection of an MDA-MB-231 primary tumor barcoded with genetic tags. From left to right: schematic overview of the dissection to isolate pieces from the center and periphery, bubble plot representing the clonal composition of each piece, and quantification of the percentage of barcodes present in each piece. Purple bars represent peripheral pieces and the orange bar represents the center. **d)** Percentage of barcodes detected in the periphery (purple) and center (orange) of tumors from different models. Each dot corresponds to a piece of primary tumor (on average, 1-2 pieces from the center and 6-12 pieces from the periphery for each tumor). Student’s unpaired t-test. MDA-MB-231 n=13, 3 independent experiments, MDA-MB-468 n= 6, one experiment, PDX-1432C n=10, 2 independent experiments, PDX-4295 n=6, one experiment, CRCM434 n=22, 4 independent experiments, CRCM412 n= 16, 4 independent experiments. **e)** Percentage of barcodes uniquely detected in the periphery (purple), tumor center (orange) or both (grey) of the tumor. Each dot corresponds to an individual tumor. Significance from one-way ANOVA followed by Tukey multiple comparisons test. **d-e)** ns: non-significant, p-value > 0.05, * = p-value <0.05, **= p-value <0.005, *** = p-value <0.0005, ****= p-value <0.0001. Error bars represent standard deviation (SD). **f)** Confocal imaging of LeGO labelled primary tumor, image of section from periphery (left) or center (middle) from MDA-MB-231 tumor, and image section from PDX-1432CC tumor, showing the center and the periphery of the tumor (right). Scale bars represent 500µm. n=1 tumor for imaging.

To investigate this further, genetic barcoding was used to study the growth of cancer clones in vivo. Breast cancer cells from the MDA-MB-231 cell line were infected with a lentivirus pool containing ∼2,600 unique genetic tags at low multiplicity of infection. Infected cells were then sorted as GFP^+^ and transplanted into the mammary fat pad of NSG (NOD/SCID/IL2Rγ^-/-^) mice, as this model and strategy were shown to produce primary tumors with multiple barcodes^40^. Tumors of 800 mm^3^ were harvested and cut into pieces of equal volume **(Fig. 1c)**. Analysis of the barcode repertoire of tumor pieces after lysis, PCR amplification, and sequencing showed that each piece was unique in its barcode composition with adjacent pieces displaying non-identical barcode repertoires, as previously described in other TNBC PDX models^9^. Interestingly, and in agreement with the simulation in **Fig. 1b**, the results confirmed that pieces corresponding to the center of the tumor (Piece #7) contained, on average, approximately 3 times more barcodes than pieces from the periphery **(Fig. 1c)**.

We then assessed whether this observation in a single tumor was reproducible across multiple mice using MDA-MB-231 and MDA-MB-468 cell lines and confirmed that this observation was consistent and reproducible **(Fig. 1d, Supp Fig. 1a-d)**. On average, the center of MDA-MB-231 and MDA-MB-468-derived primary tumors contained a large proportion of the total barcodes (66% and 77% for MDA-231 and MDA-468 respectively), compared to peripheral pieces. While the number of pieces analyzed at the center and periphery differed depending on the size and shape of the tumor, we confirmed that the higher proportion of clones in the center was not due to varying weight or volume of the tumor pieces **(Supp Fig. 1e).**

To determine whether enhanced clonal diversity in the center of the tumor could be generalized to more clinically relevant PDX models, the same experiments were performed with four different TNBC PDX models, generated from drug-naïve patient tumors **(Supp Fig. 1a)**. We confirmed that the receptor status of the original patient tumor was conserved in the PDX, based on pathology reports **(Supp Fig. 2a)**. The barcode analysis demonstrated that pieces from the center of the tumors were more clonally diverse than pieces from the periphery in PDX-4295, PDXs CRCM434 and CRCM412, recapitulating the observation made in cell line xenografts **(Fig. 1d)**. Interestingly, PDX-1432C, a highly necrotic tumor both in mouse xenografts **(Supp Fig. 2b)** and in the original patient samples **(Supp Fig. 2a)**, followed the same trend, but to a lesser extent **(Fig. 1d)**. In this case, it is possible that the necrosis resulted in the loss of many cells (and therefore barcodes) in the center of the tumor.

**Figure 2.**
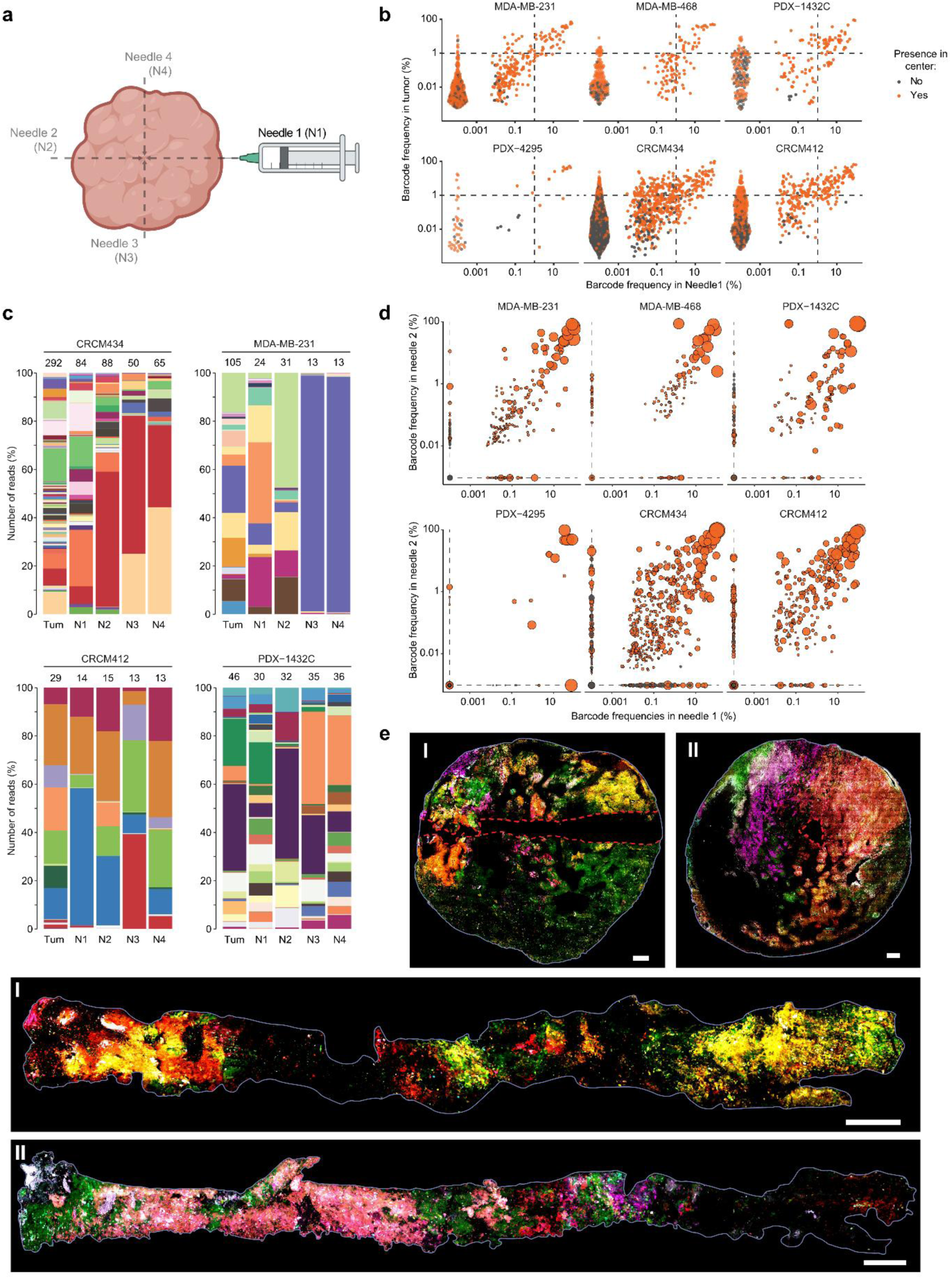
Solid biopsies: clonal composition. **a)** Schematic overview of needle sampling in a primary tumor. **b)** Clonal relationship between the frequency of barcodes detected in primary tumor and needle sample N1 for each model. Each dot represents a barcode. Barcodes present in the center are represented by orange dots and barcodes exclusively detected in the periphery are represented by grey dots. The dashed lines are drawn at 1% of barcode frequency. MDA-MB-231 n=9, 3 independent experiments, MDA-MB-468 n= 6, one experiment, PDX-1432C n=8, 2 independent experiments, PDX-4295 n=4, one experiment, CRCM434 n=19, 4 independent experiments, CRCM412 n= 16, 4 independent experiments. **c)** Examples of clonal composition of primary tumor (left column) and each needle sample (N1 to N4) for four different models represented as a stacked histogram. Each color corresponds to a barcode. The number of barcodes per sample is indicated at the top of the histogram. **d)** Relationship between barcodes detected in needle sample N2 and needle sample N1. Each dot represents a barcode, the size of the dot is correlated with the frequency of the barcode in the primary tumor. Barcodes present in the tumor center are represented by orange dots, and barcodes exclusively present the periphery are represented by grey dots. Dashed lines represent a barcode frequency of 0%. **e)** Imaging of an optically labelled primary tumor from PDX CRCM434 and of matching needle biopsies. In I, longitudinal section of the primary tumor and its needle biopsy. In II, cross section of a primary tumor and its needle biopsy. The red dashed lines highlight the needle path in the tumors, n=2. Scale bars represent 500µm.

We next determine if the localization of a clone in the primary tumor (i.e. center versus periphery) had an influence on its ability to ‘shed’ cells in distant sites **(Supp Fig. 2c)**. Interestingly, all clones detected in the lungs at high frequency were detected in both – center and periphery **(Fig. 1e, Supp Fig. 2c)**. As expected, dominant clones were present in a larger number of pieces compared to minor clones and the barcodes present exclusively in the center or the periphery were minor clones **(Supp Fig. 3a)**. Barcodes detected in the center were most often detected in several pieces from the periphery, excepted in the MDA-MB-231 xenografts **(Supp Fig. 3b, top panel)**. In contrast, barcodes detected uniquely in the periphery were often present in a low number of pieces across all models **(Supp Fig. 3b, bottom panel)**.

**Figure 3.**
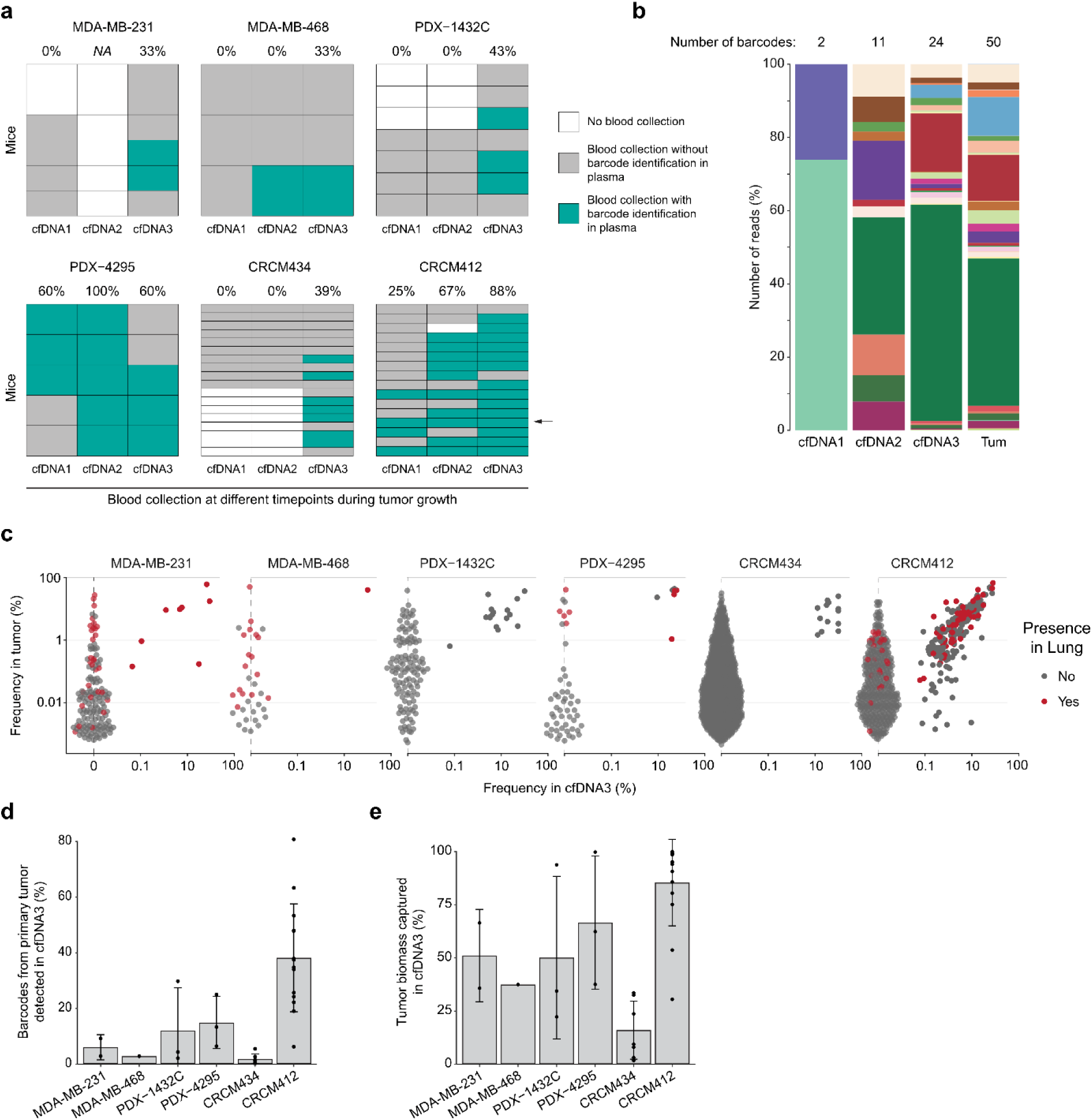
Detection of barcodes in the plasma. **a)** Barcode detection in cfDNA from early bleeding (cfDNA1 and cfDNA2) or terminal end bleed (cfDNA3). Unsuccessful barcode recovery is highlighted in grey, and successful barcode recovery in blue. The percentage of successful recovery at each time point is indicated on the top. **b)** Frequency of barcodes (based on number of reads) detected in cfDNA samples indicated with an arrow in panel **a)** and its associated primary tumor. Each color represents a barcode. **c)** Barcode frequency in primary tumors and cfDNA3. Each dot represents a barcode. The barcodes represented in red were also detected in lungs. **d)** Percentage of barcodes present in the primary tumor and detected in cfDNA3. **e)** Percentage of primary tumor biomass captured in cfDNA3. d-e) Each dot corresponds to an independent tumor. MDA-MB-231 n=2, one experiment, MDA-MB-468 n=1, one experiment, PDX-1432C n=3, 2 independent experiments, PDX-4295 n=3, one experiment, CRCM434 n=7, 3 independent experiments, CRCM412 n= 14, 4 independent experiments. The error bars represent the standard deviation (SD) of the mean.

To visualize the spatial distribution of the clones in different regions of the tumor, we used an orthogonal LeGO (Lentiviral Gene Ontology) barcoding strategy. This technology relies on a combination of fluorescent tags to visualize clones by confocal imaging^30,33,41,42^. Images from the center or the periphery of a barcoded MDA-MB-231 tumor confirmed that the center of the tumor was more clonally heterogeneous than the periphery in this model, with larger clones expanding on the ‘invasive front’ **(Fig. 1g)**. This observation corroborates a previous study that used the LeGO technology in colorectal tumors, describing that larger clones were more likely to be found in the outer region of tumors^43^. In PDX-1432C, however, the center of the tumor was highly necrotic (**Fig. 1f**, dark areas) and this result could explain why the center of the tumors barcoded with genetic tags was not as clonally dense compared to other models (**Fig. 1d**, **Fig. 1f)**.

Altogether, in silico modelling and empirical results based on genetic and optical barcoding technologies converged towards the conclusion that in preclinical models, the center of tumors is higher in clonal density compared to the periphery, that is composed of patchy clones expanding at the invasive front.

### The content of multiple needle biopsies from a given tumor are highly variable, but likely to contain dominant clones

Solid biopsies are routinely used for the diagnosis and prognosis of breast cancer patients. As our results in barcoded models confirmed the uneven distribution of the clones across the tumor previously observed in PDXs^9^ and patient samples^18,20,35^, we interrogated the level of ITH captured using needle sampling in these barcoded models. To do so, multiple needle samplings were taken from primary tumors **(Fig. 2a)**. Needles were directed towards the center of the tumor in 4 different directions and the barcode repertoire captured in the ex vivo biopsies was analyzed and compared to the repertoire from the whole tumor. In general, and as expected, the barcode frequency detected in one needle correlated with the barcode frequency in the primary tumors **(Fig. 2b)**. However, several dominant clones from the primary tumors were not captured in the biopsy.

When comparing multiple biopsies from the same tumors, we found that the barcode repertoire varied depending on the orientation of the needle **(Fig. 2c)**, with several minor and dominant barcodes not shared between biopsies collected in opposite directions **(Fig. 2d, Supp Fig. 4a, 4b)**. This was consistent with earlier findings that clones grow as patches in specific sub-regions of the tumor **(Fig. 1f)** and that the composition of solid biopsy is largely influenced by the distribution of the clones in the tumor **(Fig. 2e)**.

**Figure 4.**
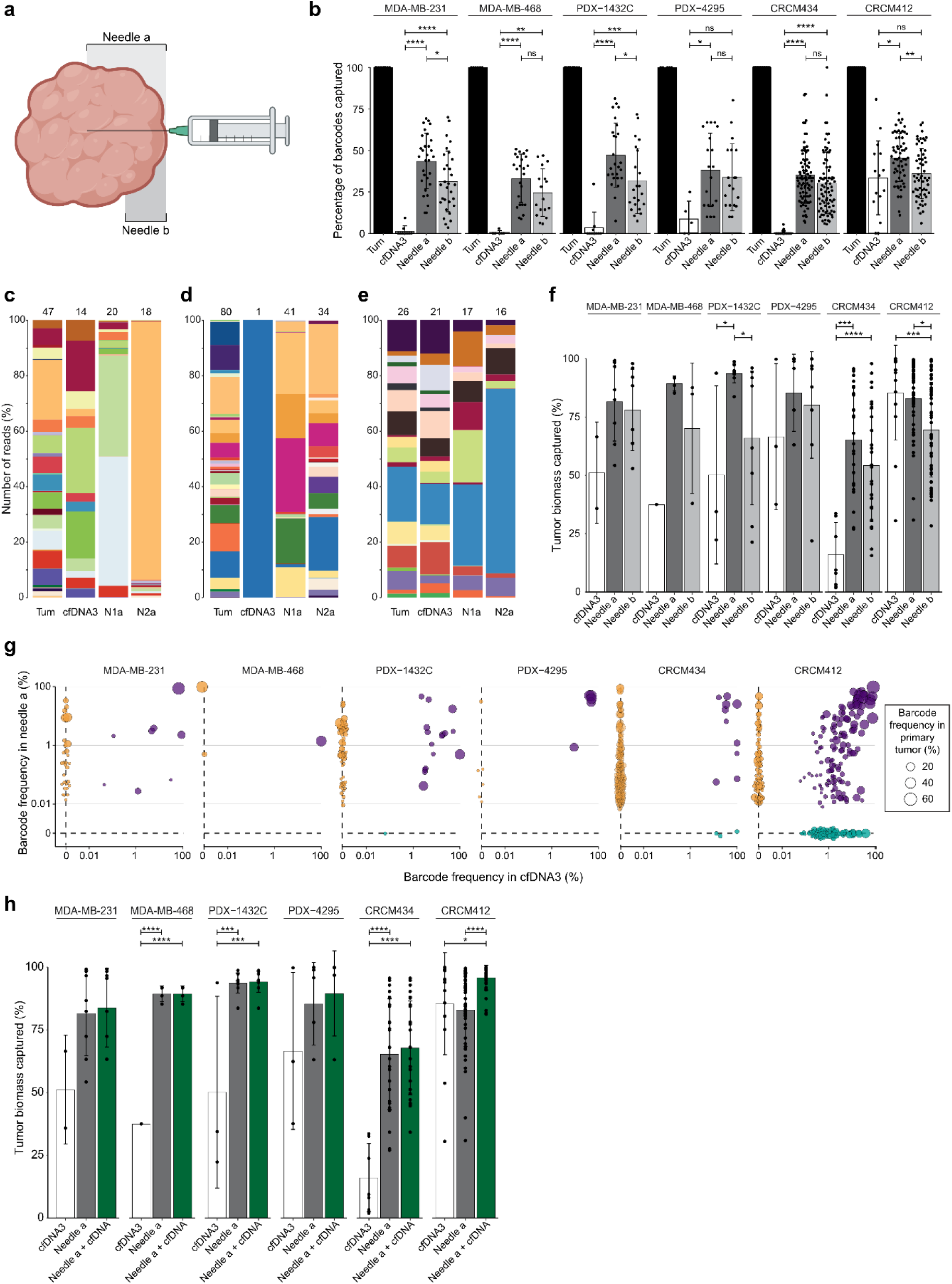
Comparison of the barcode repertoire captured in liquid and solid biopsies. **a)** Schematic overview of needle depth during sampling in a primary tumor. **b)** Percentage of barcodes detected in needle biopsies from the primary tumor (plotted as reference at 100%) and cfDNA. In this panel, all the cfDNA samples were included, including those with unsuccessful barcode recovery. Needle a corresponds to solid biopsy reaching the center of the tumor (deep needle sampling). Needle b corresponds to shallow sampling, covering only a quarter of the tumor diameter. Each dot corresponds to a biopsy sample. **c-e)** Three representative examples of the clonal repertoire detected in primary tumors, cfDNA3 and two needles sampling in the PDX model CRCM412. In this stacked histogram, each barcode is represented by a color. **f)** Percentage of the primary tumor biomass detected in different biopsies. **g)** Relationship between barcodes in solid biopsy (Needle 1) and cfDNA (cfDNA3), depending on the barcode frequency in primary tumor (represented by the size of the dots). Each dot represents a barcode. Barcodes detected within the two biopsies are highlighted in purple and barcodes exclusively detected by a method are plotted at 0, by needle (orange) and in cfDNA (cblue). **h)** Primary tumor biomass captured in different biopsies: cfDNA (white), needles (grey) and combination of deep needle sampling (Na) and cfDNA3 (green). **b,f,h)** Significance from one-way ANOVA followed by Tukey multiple comparisons test, * = p-value <0.05, **= p-value <0.005, *** = p-value <0.0005, ****= p-value <0.0001. Error bars represent standard deviation (SD). **f,h)** Each dot corresponds to a biopsy sample. In these panels, only samples in which barcodes were detected in the plasma were included in these panels. MDA-MB-231 n=2, one experiment, MDA-MB-468 n=1, one experiment, PDX-1432C n=3, 2 independent experiments, PDX-4295 n=3, one experiment, CRCM434 n=7, 3 independent experiments, CRCM412 n= 14, 4 independent experiments.

While a dominant barcode in a biopsy was likely to be a dominant barcode within the primary tumor **(Fig. 2b and Fig. 2c)**, some dominant clones in the tumor were missed in the biopsy. Overall, a needle biopsy that reached the center of the tumor rarely reflected the full barcode heterogeneity of the primary tumor. However, these biopsies contained clones that were representative of 80 to 90% of the total biomass of the tumor **(Supp Fig. 4c)**.

### Barcode detection in cfDNA depends on the tumor model and tumor burden

While the clonal repertoire captured in solid biopsies was biased by the spatial distribution of the clones within the tumor, we hypothesized that liquid biopsies, and in particular cfDNA isolated from the blood, may overall reflect the clonal makeup of the primary tumor.

To test this hypothesis, we assessed the presence of the genetic barcodes in the cfDNA of tumor bearing animals. We collected blood via the tail vein at different time points, when tumors reached 100 mm^3^ (cfDNA1), 300 mm^3^ (cfDNA2), or via heart terminal end bleed when the tumors reached 800 mm^3^ (cfDNA3). After plasma separation, DNA was extracted and barcodes amplified by PCR run in five-replicates to increase the probability of barcode recovery. The results indicated that barcodes were not consistently detected in all mice from the same cohort **(Fig. 3a)**. At the last time point (cfDNA3), the larger volume of blood collected and the type of bleeding (heart bleed rather than tail vein, see methods) could contribute to the increase of barcode detected. However, the overall barcode recovery seemed to increase with tumor burden, as shown when comparing cfDNA1 and 2 in the MDA-MB-468 and CRCM412 bearing mice **(Fig. 3a and b)**.

Interestingly, the number of barcodes detected in the blood largely depended on the tumor model. For example, PDX-4292 and CRCM412 had a higher level of barcode recovery in plasma, even when tumors were small **(Fig. 3a)**. This suggests that the onset of cfDNA shedding might vary between tumors, depending on the intrinsic properties of the clones.

Where possible, we compared the identity of the barcodes detected in the plasma over time **(Fig. 3b)**. Surprisingly, we found that different barcodes were detected at different times in the same mice. The barcodes detected in cfDNA at early bleeding didn’t necessarily reflect the ITH of the primary tumor analyzed at ethical endpoint. On the contrary, cfDNA collected at the time of primary tumor harvest (cfDNA3) contained a barcode repertoire that was representative of the barcode repertoire of the primary tumor **(Fig. 3b, 3c, Supp Fig. 5a)**. Even though recovery of barcodes from cfDNA greatly differed between models **(Fig. 3a)**, when detected, only a small percentage of the barcodes from the primary tumor were represented, except for PDX CRCM412 **(Fig. 3d, Supp Fig. 5b)**, and some of the dominant barcodes in the primary tumors were not detected in the plasma **(Fig. 3b, 3c)**. However, the clones detected in cfDNA were contributing a significant percentage of the tumor biomass **(Fig. 3e, Supp Fig.5b,** up to ∼80% for PDX CRCM412), representing clones present in the center and periphery of the tumors **(Supp Fig. 5c)**. Furthermore, cfDNA seemed to be a good surrogate of the heterogeneity captured in lungs as highlighted in **Fig. 3c**. Overall, the barcodes captured in cfDNA and needle biopsies represented a similar proportion of the tumor biomass detected in lungs **(Supp Fig. 5d)**.

As the barcode repertoire detected in blood varied over time **(Fig. 3b)**, we investigated whether this result was due to differences in the clonal composition of the primary tumor. To do so, the same clones from PDX-1432C were implanted into multiple mice, after an *in vivo* expansion of the clones. These “sister” mice were then harvested when the primary tumors reach different sizes, and the same volume of blood was collected for each mouse. The barcode repertoire was analyzed in primary tumor and plasma **(Supp Fig. 5e)**. When the barcode recovery was compared in the different mice of these cohorts, we confirmed that, in agreement with the results presented in **Fig. 3a**, the recovery rate of barcodes in the plasma depended on the size of the primary tumor **(Supp Fig. 5e)**.

The clonal repertoire of tumors between sister mice from each experiment was strikingly conserved over time **(Supp Fig. 5f)**. However, the barcode recovery from cfDNA was variable between mice and barcode frequency identified slightly differed from one mouse to the other **(Supp Fig. 5f)**. These differences could be due to the low level of cfDNA detected in these models, or due to variations in cfDNA shedding depending on tumor vascularization or the spatio-temporal evolution of the clones in individual mice.

### cfDNA and solid biopsies can provide complementary information

As both solid and liquid biopsies captured varying degrees of ITH, we compared the clonal repertoire using each of these sampling techniques on the same mice across the 6 xenograft models. Overall, taking into consideration the cases where cfDNA was not detected, we found that solid biopsies reaching the center of the tumor or half of the radius, captured a larger percentage of the primary tumor barcodes compared to cfDNA **(Fig. 4a, b)**. However, for those mice with barcodes detected in the plasma, the Pearson correlation between needle sampling or cfDNA and primary tumor did not show any significant difference **(Supp Fig. 6a, 6b)**. It is important to note that in these pre-clinical models, the barcode content of biopsies varied significantly, not only between models, but also within a model, regardless of the method. This could be due to the low detection rate and resulting ‘randomness’ obtained when sampling small amount of tissues and limited blood volumes. Indeed, comparing the identity of barcodes captured by the two approaches in the same animal **(Fig. 4c-e)**, three distinct patterns were identified. In the first **(Fig. 4c),** the barcode repertoire from each sample was unique, each needle biopsies and liquid biopsies containing different sets of barcodes. In this case, the cfDNA barcode repertoire better recapitulated tumor heterogeneity. In the second **(Fig. 4d),** solid biopsies were a better surrogate of primary tumor ITH, while cfDNA only captured one clone from the primary tumor. The last pattern **(Fig. 4e)** showed similarity between cfDNA and needle biopsies, where both biopsy methods accurately represented the primary tumor.

The tumor biomass captured with each method was then quantified **(Fig. 4f)**. Needle sampling was a better compendium of primary tumor heterogeneity compared to cfDNA (when detected in plasma), except for CRCM412 where cfDNA overall captured more tumor biomass than the two needle methods.

We then assessed whether the same clones were captured with both methods and whether any advantage in the assessment of ITH was conferred by combining several biopsies. Barcodes captured by both cfDNA and solid biopsies were often similar **(Fig. 4g, orange circles)**, while solid biopsies seemed to have more unique barcodes **(Fig. 4g, black circles on the y-axis)** and were more efficient at capturing minor barcodes **(Supp Fig. 6b)** in all models except CRCM412. When two needle samples were combined from the same primary tumor, the overall primary tumor biomass captured from the primary tumor slightly increased **(Supp Fig. 6c)**. Furthermore, combining cfDNA and needle biopsies significantly increased the percentage of biomass captured compared to cfDNA alone **(Fig. 4h, Supp Fig. 6d)**, with a few exceptions. In addition, the combination of solid and liquid biopsies significantly increased the percentage of biomass captured in CRCM412 **(Fig. 4h)**. However, it didn’t provide any significant advantage in the other models **(Fig. 4h, Supp Fig. 6d, Supp Fig. 7a)**.

Lastly, in the MDA-MB-231 model, CTCs were successfully retrieved from blood, and the barcode repertoire was analyzed **(Supp Fig. 7b)**. While barcodes detected when combining needle biopsies and cfDNA already represented a high percentage of the primary tumor biomass, adding the barcodes collected in CTCs further improved the assessment of ITH, covering over 80% of tumor biomass **(Supp Fig. 7b)**.

## Discussion

In clinical settings, probing tumor heterogeneity is of utmost importance for the ongoing development of precision medicine. As drug resistance and metastasis can be driven by minor clones present in primary or secondary lesions^44^, there is a need to capture molecular information from a large number of clones, regardless of their frequency, to guide the choice of optimal therapies. Cellular barcoding was used here to generate clonally structured tumors in mice, offering a powerful qualitative and quantitative assessment of the extent of ITH captured in solid and liquid biopsies in two breast cancer cell lines and four PDXs. Overall, the superiority of one sampling method over another clearly depends on the patient model and tumor burden. This suggests that, in the clinic, combining tumor and blood samplings could provide a better assessment of ITH.

Our results suggest that solid biopsies can capture a large range of minor and dominant clones present in primary tumors across models, representing ∼60-90% of the primary tumor biomass. However, the clonal repertoire captured in solid biopsies is strongly biased by the spatial distribution of the clones in the primary tumors. As a result, multiple sampling of a tumor is likely to provide highly variable results in heterogeneous tumors. While it would be interesting to determine whether this cellular heterogeneity (based on barcode identification) correlates with genomic heterogeneity, this result corroborates previous observations based on genomic analysis of serial biopsies^45^. Regional sequencing of patient tumors^18,20,36–38^ and recent special transcriptomic analysis^35,46^ also support the observation that tumor sampling is not necessarily representative of the whole tumor heterogeneity.

In the clinic, needles are often oriented towards the center of a tumor. Indeed, both computational modeling and empirical data from cellular barcoding experiments highlighted that the center of non-necrotic tumors is significantly enriched in barcodes compared to the periphery. While a similar observation has been made in colorectal xenografted tumors^43^, it would be important to confirm this in immunocompetent preclinical models and patient samples. This observation is extremely timely, as many studies are currently investigating the transcriptomic profile of cancer clones using spatial transcriptomic studies^35,46–48^. It will be important to determine whether this difference in clonal frequency and distribution can be attributed to particular features of the clones. It might be that cells from each subclone have different gene expression profiles depending on their localization within the tumor. Breast cancer stem cells from the center, for instance, were shown to be more epithelial than cancer stem cells present in the invasive front^49^. Further analysis using spatial transcriptomic analysis, optical barcoding and time-lapse imaging may allow the study of these mechanisms over time.

In parallel, quantitative comparison between barcodes in mouse plasma and barcodes in whole tumors suggested that when detected, cfDNA can a good surrogate of tumor heterogeneity as it allowed the capture of up to 80% of the tumor biomass. However, the likelihood of detecting barcodes in cfDNA correlated with the extent of tumor burden, and dominant barcodes present in primary tumors and lungs had a higher likelihood to be detected in cfDNA, corroborating previous patient studies^13,24,50,51^. It is also possible that necrotic models (such as PDX-1432C) were more likely to shed DNA into the vasculature compared to non-necrotic models (such as MDA-MB-231 xenografts). Nonetheless, discrepancies were identified between models, and tumor burden and extent of necrosis were not solely accountable for these differences. In the case of PDX CRCM412, for instance, cfDNA can be detected at early time points, despite its low-necrotic grade. Interestingly, cells were detected in the lungs in this model, and it might be that tumors able to shed cancer cells in their bloodstream are also more likely to shed cfDNA. It would be interesting to validate this hypothesis and determine whether the presence of previously described ‘shedders’ and ‘seeders’ correlates with the presence of cfDNA in the plasma. Understanding the mechanisms involved in cfDNA shedding will be clinically relevant.

cfDNA has a promising role in the quantitative monitoring of early recurrence and tumor progression, and it also has qualitative utility as a molecular profiling tool. Tests such as FoundationOne Liquid CDx have been FDA approved to guide therapeutic interventions in several types of cancers (non-small cell lung cancer, breast, prostate and ovarian cancers) based on a panel of somatic and germline mutations. Therefore, understanding the phenotype of the clones that are likely to be detected in the plasma will be useful in the interpretation of liquid biopsies. While our results indicate that barcodes from cfDNA predominantly represented dominant clones from the primary tumors, it would be interesting to determine whether these clones have seeding properties^9^ and might, therefore, be responsible for the establishment of metastases. To study the ability of cfDNA to reflect the heterogeneity of metastases, it would be interesting to compare the barcode repertoire detected in plasma compared to metastases after resection of the primary tumor. In this case, cfDNA may be a good surrogate of the biomass of metastases from different sites that are difficult to sample with solid biopsies.

The qualitative analysis of barcodes in biopsies suggested that barcodes detected in cfDNA were likely to represent dominant clones and may fail at capturing clones that were under-represented in the primary tumor, which may be responsible for distal recurrence^9,29^ and drug resistance^44^. However, these results, based on the detection of genetic barcodes that are smaller than 100bp, may underestimate the representation of cfDNA compared to 100bp fragments from the whole genome detected in clinical settings^13^. It is possible that some barcodes present in truncated forms were not recognized for primer annealing and therefore not detected. Furthermore, the volume of blood and bleeding strategy used in these pre-clinical models differed from clinical settings.

While this quantitative analysis in PDX might not be directly comparable to analysis in the clinic, it provides an opportunity to study the process of DNA shedding by cancer clones in a longitudinal manner, in the absence of therapeutic interventions. Interestingly, variabilities in liquid biopsies were observed in mice bearing tumors with similar clonal composition, highlighting that some of these variations might be stochastic. Further studies linking genomic analysis and clonal information in these models will be required to better understand the process of DNA shedding.

Finally, our results demonstrated that combining liquid and solid biopsies can provide a significant advantage in the assessment of ITH, depending on the tumor model. This observation supports other studies indicating that it might be difficult to substitute one type of biopsy with the other^24,52^, and that ensuring that both techniques coexist would improve breast cancer diagnosis and management^23,53^. Indeed, sequencing analysis of 351 samples from patients with diverse cancer types suggested that the combination of both solid and liquid biopsies offers a more therapeutically valuable representation of tumor heterogeneity in clinical settings^23^. Furthermore, solid biopsies present the advantage of capturing intact cells, increasing the scope of downstream applications in diagnostics and research from cellular assays to studying malignant or normal cells^54,55^, to multi-omics analysis based on bulk or single cells^8,56^. From a diagnosis and prognosis point of view, they provide a unique platform to analyze clinically relevant markers that are complementary to the study of cfDNA, such as the presence of tumor infiltrating lymphocytes predictive of immunotherapy efficacy.

While more work will be required to further understand the properties and dynamics of cancer clones in different types of biopsies over time, and in response to specific therapies, this study provides new insights into the utility of barcoded models to study the variability and biases of solid and liquid biopsies. Linking meaningful clonal information to multi-omics analyses in pre-clinical models of cancer holds great promise in the development of diagnostic tools and personalized therapies.

## Material and Methods

### *In silico* modeling of 3D growth

Simulations of 3D growth of barcoded tumour cells were performed as previously described^9^. In brief the simulation code by Waclaw, B. et al^39^ was adapted to account for cellular barcoding and transplantation into the mammary fat pad, using the parameters originally proposed by the authors for growth and migration. Simulations were initiated with 200 barcoded tumour-initiating cells and ran until reaching a size of 10 million cells. To quantify clonal density, the simulated tumour was split *in silico* into 5 pieces (as shown in Fig 1b), and the number of clones counted, and their frequencies plotted. The positions of each cell as well as clonal identity (color) were rendered in 3D using the open source visualization tool Ovito^57^.

### PDX establishment and amplification

PDX-1432C was established at ONJCRI, from a triple-negative treatment naïve breast cancer tumor. KCC-P-4295, referred to in the manuscript as PDX-4295, was obtained from BROCADE and established at the Kinghorn Cancer Centre and Garvan Institute, from a treatment naïve TNBC. PDXs CRCM434 and CRCM412 were generated at the Institut Paoli-Calmettes from drug naïve TNBC patient tumors. All PDXs were orthotopically injected into the mammary fat pad of females NOD-SCID-IL2Rγ -/- (NSG) to be amplified prior to barcoding. All procedures in animals were conducted in accordance with the National Health and Medical Research Council guidelines under the approval of the Austin Animal Ethics Committee. The use of patient samples was approved by Austin Health Human Research Ethics Committee.

PDXs tumors were harvested and prepared as a single-cell suspension. The tissues were manually chopped into small pieces (about 1 mm by 1 mm) and resuspended for one hour in the following digestion medium: collagenase IA (300 U/ml) (#C9891, Sigma-Aldrich), hyaluronidase (100 U/ml) (#H3506, Sigma-Aldrich), and deoxyribonuclease I (DNase I) (100 U/ml) (#LS002139, Worthington) in DMEM/F12 (#10565042, ThermoFisher. PDXs cells were plated in 24 well plates (flat bottom ultralow attachment, #734-1584, Corning) at a density of 300,000 cells in 300ul of mammosphere media. Mammosphere medium was composed of DMEM-F12 (#10565042, ThermoFisher) supplemented with 1X B27 (#17504001 ThermoFisher), 100U/ml of penicillin-streptomycin (#15140122, ThermoFisher), 5ug/ml insulin (#11376497001, Sigma-Aldrich), 1ug/ml hydrocortisone (#H0396-100MG, Sigma-Aldrich), 0.8U/ml heparin (#H0878-100KcU, Sigma-Aldrich), 20ng/ml basic fibroblast growth factor (#01-106, Merck-Millipore), and 20ng/ml epidermal growth factor (#E9644, Sigma-Aldrich).

### Optical barcoding experiments using the LeGO system

Lentiviruses were produced by transient cotransfection of human embryonic kidney–293T cells with the LeGO vectors of interest and the packaging plasmids using a FuGENE HD transfection reagent (#E2311, Promega) following the manufacturer’s recommendations. LeGO-EBFP2 (Addgene plasmid # 85213 ; http://n2t.net/addgene:85213 ; RRID:Addgene_85213); LeGO-S2 (Addgene plasmid # 85211 ; http://n2t.net/addgene:85211 ; RRID:Addgene_85211); LeGO-V2 (Addgene plasmid # 27340 ; http://n2t.net/addgene:27340 ; RRID:Addgene_27340); LeGO-T2 (Addgene plasmid # 27342 ; http://n2t.net/addgene:27342 ; RRID:Addgene_27342); LeGO-dKatushka2 (Addgene plasmid # 85214 ; http://n2t.net/addgene:85214 ; RRID:Addgene_85214); LeGO-C2 (Addgene plasmid # 27339 ; http://n2t.net/addgene:27339 ; RRID:Addgene_27339); LeGO-mOrange2 (Addgene plasmid # 85212 ; http://n2t.net/addgene:85212 ; RRID:Addgene_85212) were a gift from Boris Fehse^41,42^. pCMVR8.74 (Addgene plasmid # 22036 ; http://n2t.net/addgene:22036 ; RRID:Addgene_22036) and pMD2.G (Addgene plasmid # 12259 ; http://n2t.net/addgene:12259 ; RRID:Addgene_12259) were a gift from Didier Trono. The virus production and infection of cells of interest were generated as previously described^30^. In brief, PDXs and cell line MDA-MB-231 were prepared and maintained as described above. After plating, the cells were kept in culture for 72 hours in the presence of a combination of LeGO-Viruses. For PDX CRCM434 and MDA-MB-231, 5 fluorescent proteins were used: eBFP2, tSapphire, Venus, tdTomato and dKatushka2. For PDX 1432C, a combination of 6 fluorescent proteins was used: eBFP2, tSapphire, Venus, mCherry, mOrange and dKatushka2. Infected cells were then sorted before transplantation into NSG mice.

Harvested tumors were fixed in freshly made 4% paraformaldehyde (#28908, ThermoFisher) in DPBS. Tumors were kept in fixation buffer for 24h at 4°C on a roller protected from direct light. Samples were then washed with DPBS and embedded in 3% low-melting point agarose (#1613111, Bio-Rad). For the needle samples, a 23G needle was inserted in the primary tumor and the resulting biopsy was imaged as described below. 200µm sections were cut and mounted on slides with mounting medium (#P36961, ThermoFisher) and high-performance coverslips (#474030-9000-000, Zeiss Microscopy). Image acquisition was performed on a confocal microscope (LSM 880, LSM 980, Zeiss).

### Genetic barcoding experiments

For PDXs, cells were infected with lentiviruses containing the barcode library as previously described^9^, at low MOI (PDX-1432CC 8.7±1.1%, CRCM434 6.4±2.9%, PDX-4295 7.5±3.5%, CRCM412 5±1.5% (mean±SD)) to ensure the integration of a single barcode per cell. Barcoded cells were sorted for GFP positivity and resuspended in injection buffer (42.5% DPBS, 30% FBS 25% Matrigel and 2.5% Trypan blue). 2.5-5k of cells were injected into the fourth mammary gland of NSG mice. For PDX CRCM412, the barcode composition of primary tumors and metastases from mice 88-92 have been previously described^40^.

To generate the models used for cloning experiments, barcoded tumors were harvested once they reached 300mm^3^, processed into single-cell suspension, and reinjected in 12 recipient mice.

For the cell lines, cells were obtained from the ATCC, MDA-MB-231 (#HTB-26, passage 39), and MBA-MB-468 (#HTB, passage 339) were maintained in RPMI 1640 with Hepes (#22400086, ThermoFisher), 10% fetal bovine serum (FBS) and penicillin-streptomycin (#15140122, ThermoFisher) at 10,000U/ml.

For barcode infection, 2.2 million cells were plated in 10cm cell culture dishes (#353003, Corning) with 8ml culture media. Lentiviruses were added for 48h hours. Barcoded cells were selected via flow cytometry based on GPF positivity, both populations under 0.1 MOI, 7.1% and 5% for MDA-MB-231 and MDA-MB-468 respectively, 25,000 cells per cell line were then expanded in vitro. 200k MDA-MB-231 barcoded cells and 300k MDA-MB-468 cells were injected into the mammary fat pad of NSG mice. Mice 36-39, 69-72 were previously analyzed for barcode composition in primary tumors and metastases^40^.

### Tumor processing and needle biopsies

Genetically barcoded tumors were harvested once they reach 800mm^3^, prior to needle sampling. A preloaded 3ml syringe with 500ul DPBS fitted with a 23G needle was inserted in the primary tumor with minimal aspiration to ensure the capture of tissue. The biopsy content was collected in a 1.5μl Eppendorf tube. Needles and syringes were changed between each biopsy. Tubes containing biopsy samples were spun down 5min at 500rpm, and the supernatant was removed. The pellets were resuspended in 50ul lysis buffer (Viagen Biotech) with 1:50 proteinase K 20mg/mL (Invitrogen) and lysed 1h at 55°C followed by 30min at 85°C, and finally 5min at 95C on a heater shaker dry bath set to 800rpm. Samples were stored in a freezer until PCR amplification.

Tumor dissections were performed with surgical blade n°10 in order to isolate tumor center from edges as shown in **Fig. 1c**. Edges on the transversal axes were cut first from the primary tumor, then the lateral edges were removed from the primary tumor. The tumor was then flipped horizontally to dissect superior and inferior edges, leaving the center of the primary tumor exposed. Pieces were re-cut if necessary to obtain pieces of equal size. Blades were changed between each tumor to avoid barcode cross-contamination. Pieces were resuspended in 300μl of lysis buffer and lysed overnight on heater shaker set to 800rpm at 55°C then 85°C 30min and 95°C for 5min. Samples were stored in a freezer until PCR amplification.

### Blood collection and cfDNA isolation

Blood from mice was collected to isolate cfDNA. Terminal end bleed was performed via cardiac puncture after death with a 1ml 27G insulin syringe, 800µl to 1ml of blood was collected in microvette tubes (Sarstedt). For early time points, 200µl of blood was collected via the lateral tail vein with a 0.5ml 27G insulin syringe and transferred to a heparin coated tube. Blood samples were immediately processed. Blood tubes were spun down at 1600g for 10min in a “swing-out” rotor centrifuge set with the brakes off Plasma was transferred to 1.5ml tubes and centrifuged for 5min at 14000rpm to pellet cellular debris. The supernatant was transferred into a new tube and stored at -80°C until cfDNA extraction. cfDNA extraction was performed using the QIAamp Circulating Nucleic Acid kit (Qiagen).

### Lung metastasis analysis

Lungs were collected at ethical endpoint. The tissue was manually chopped, and the digestion was done in 5ml of RPMI 1640 for MDA-MB-231 and MDA-MB-468 cells lines, and DMEM F12 (ThermoFisher) for PDX cells. Both media were supplemented with 300U/ml collagenase IA (Sigma Aldrich), 100U/ml hyaluronidase (Sigma Aldrich). Samples were incubated at 37°C on an orbital shaker set at 300rpm for 45min, then resuspended through a 18G needle after 20min, and a 21G needle after 40min of digestion. The cell suspension was filtered through a 70µm cell strainer and spun down for 5min at 500g. PDX samples were resuspended in DPBS with PI to be sorted via flow cytometry. Sorted cells were spun down and resuspended in 50µl lysis buffer. Cells from the cell lines were resuspended in 100µl lysis buffer. Samples were lysed for 1h at 55°C followed by 30min at 85°C, and finally for 5min at 95°C on a heater shaker set to 800rpm. Samples were stored in a freezer until PCR amplification.

### Barcode amplification and sequencing

PCR amplification was performed on crude lysates, tumor pieces were diluted 1:10 in water. 40µl of this template was mixed with 160µl of PCR mixed in a well of 96 well-plate (#3420-00S, SSIbio) and split in two replicates of 100µl before the start of the PCR to assess barcode detection reliability. cfDNA and lung samples were run in quintuplicate. The first PCR included common primers (TopLib 5’-TGCTGCCGTCAACTAGAACA-3’ and BotLib 5’-GATCTCGAATCAGGCGCTTA-3’) to allow for barcode amplification. Cycle specification was at 94°C for 5min, followed by 30 cycles at 94°C for 15sec, 57.2°C for 15sec, 72°C for 15sec, and then 72°C for 10min. Product of the first PCR was then used to run the second PCR, to add specific individual indexes for NexGen sequencing. Cycle specifications of the thermocycler were 94°C for 5min, followed by 30 cycles at 94°C for 5sec, 57.2°C for 5sec, 72°C for 5sec, and then 72°C for 10min. All PCR were run on abS100 thermal cycler (Bio-Rad), and the final presence of PCR product at 266bp was verified on 2.5% agarose gel electrophoresis. Samples were pooled and clean-up with magnetic beads (#744100.4, Macherey-Nagel) before sequencing on Next-Seq (Illumina).

### Histology

Tissues were fixed in 4% paraformaldehyde for 48h before transfer to 70% Ethanol, for block embedding, and staining was performed by the department of pathology at the Austin Health Hospital. Slides were scanned on Aperio AT2.

### Bioinformatic analysis

Sequencing results were analyzed on RStudio (Version 1.4.1106), demultiplexing of sequencing FastQ files were performed using the ProcessAmplicon function from the edgeR package (DOI: <u>10.18129/B9.bioc.edgeR</u>) to generate a read-count matrix for each barcode per sample. To ensure the quality of the data, multiple filtering was applied to the data set. Once separated by sample, barcode read counts below or equal to 10 were set to zero, then replicate and quintuplicate samples with a Pearson correlation inferior to 0.6 were removed from the analysis (except for cfDNA samples). Finally, barcodes present in less than 2 replicates were discarded. Replicates were then pooled by adding read count values of the barcode and normalized. The value of each barcode within individual tumor pieces were cumulatively added and normalized to recreate the profile of the full tumor as displayed in **Figure 2C**. All subsequent visualization and statistical analysis were performed in R.

## Data and Code availability

The data that support the findings of this study and the code to reproduce all relevant figures and supplementary figures included in this article are available here: https://github.com/Anto-Ser/Biopsies_Barcoding.git.

## Acknowledgements

We would like to thank Robin Anderson, Bhupinder Pal, Caroline Bell, Sarah Ellis, David Baloyan and Stephen Wilcox for their advice and technical assistance. The authors and Olivia Newton-John Cancer Research Institute gratefully acknowledges the generous support of the Love Your Sister Foundation. We are grateful to the patients who consented for their tissue to be donated. We are also grateful for the support of the BROCADE Rapid Autopsy Program, funded by the Australian National Breast Cancer Foundation (NBCF) under infrastructure grant IF-14-001, to facilitate access to PDX-4295. We also acknowledge Lisa Devereux, Robin Anderson and Alex Swarbrick, who manage and lead BROCADE. We acknowledge the Kinghorn Cancer Centre and Garvan Institute for the KCC-P-4295 model (PDX-4295), the Cancer Research Center of Marseille (CRCM) for PDXs CRCM412 and CRCM434, and Ton Schumacher and the Netherlands Cancer Institute for providing the genetic barcoding library. The Olivia Newton-John Cancer Research Institute acknowledges the support of the Operational Infrastructure Program of the Victorian Government. AS was supported by the Melbourne Research Scholarship, DM is supported by the Victorian Cancer agency (MCRF21011), the NBCF (Investigator Initiated Research Grant IIRS0049) and NHMRC (GNT2012196 and 2027459). DM and BY are supported by the Love Your Sister Foundation. The authors acknowledge the ACRF Centre for Imaging the Tumour Environment at the Olivia Newton-John Cancer Research Institute for providing microscopy support, and the Austin Pathology.

## Author contributions

AS, TSW, SJD, SN and DM designed the experiments and wrote the manuscript, AS performed the genetic barcoding experiments, AS and TSW performed the computational and statistical work, JB, VCW and KLR performed the LeGO experiments, SF performed the cfDNA isolation, JB and FES helped with the *in vivo* experiments, EL provided PDX-1432C and PDX-4295, ECJ and CG provided CRCM412 and CRCM434, DW, FH and BY contributed expertise and resources, all authors discussed the results and provided some feedback on the manuscript.

## Supplementary figures

**Supplementary Figure 1:**
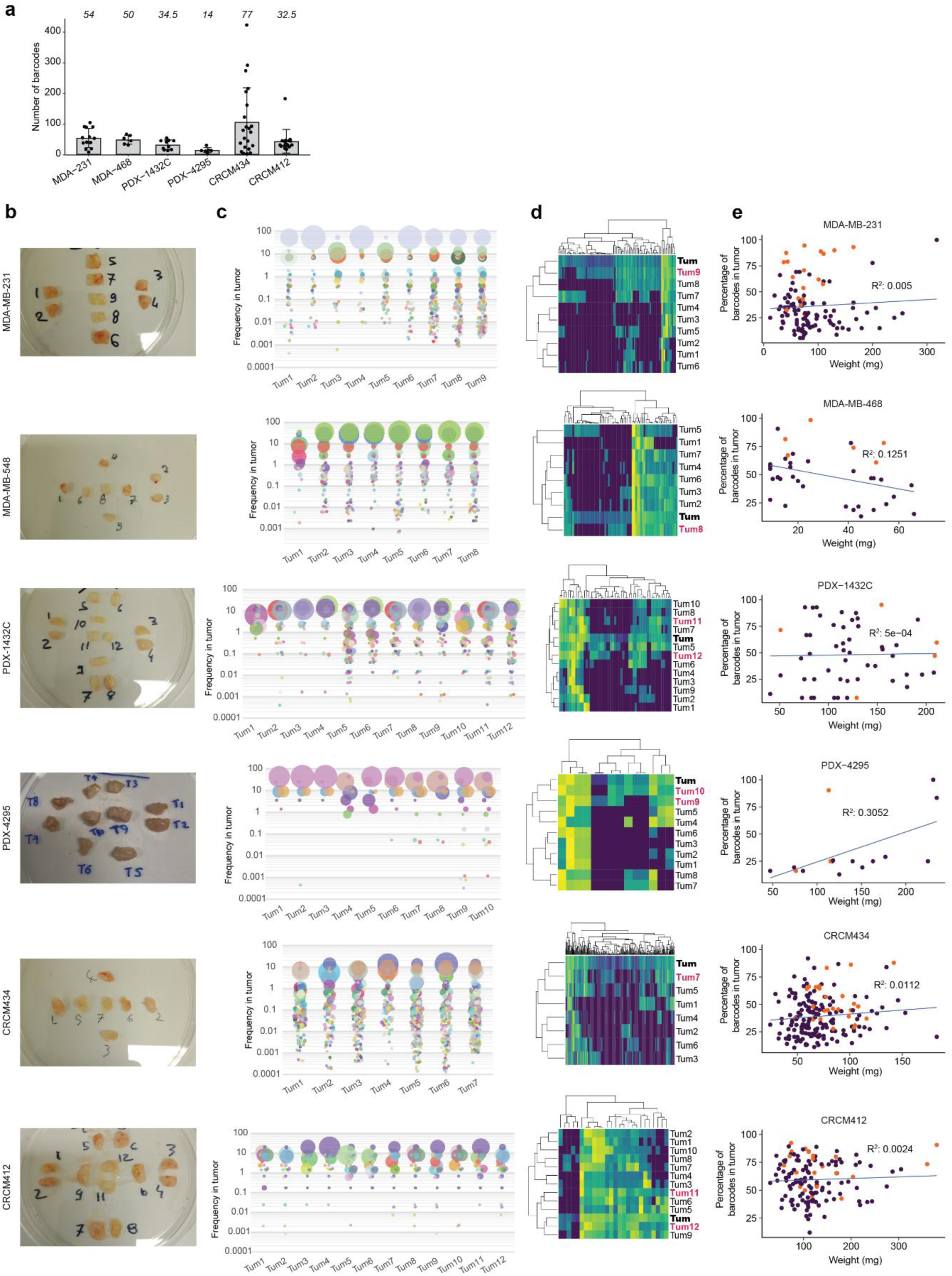
**a)** Total number of barcodes detected in each tumor, MDA-MB-231 n=13, 3 independent experiments, MDA-MB-468 n= 6, one experiment, PDX-1432C n=10, 2 independent experiments, PDX-4295 n=6, one experiment, CRCM434 n=22, 4 independent experiments, CRCM412 n= 16, 4 independent experiments. Each dot correspond to a tumor. Error bars represent standard deviation (SD). **b-e)** Example of primary tumor cutting and barcode composition for each model, from top to bottom, MDA-MB-231, CRCM434, PDX-4295, PDX-1432C, CRCM412, MDA-MB-468. **b)** Tumors were collected in PBS and cut into pieces of similar size. **c)** Representation of the clonal composition of each piece. **d)** Heatmap representing the clonal composition of each peripheral piece (normal font), center pieces (red) and full tumor (bold). Each column represents a barcode, and its frequency is represented by the color scale. **e)** Relationship between the percentage of barcodes detected in each tumor piece and the weight of the tumor piece. Each dot corresponds to a tumor piece, from the periphery (purple dots) or the center (orange dots).

**Supplementary Figure 2:**
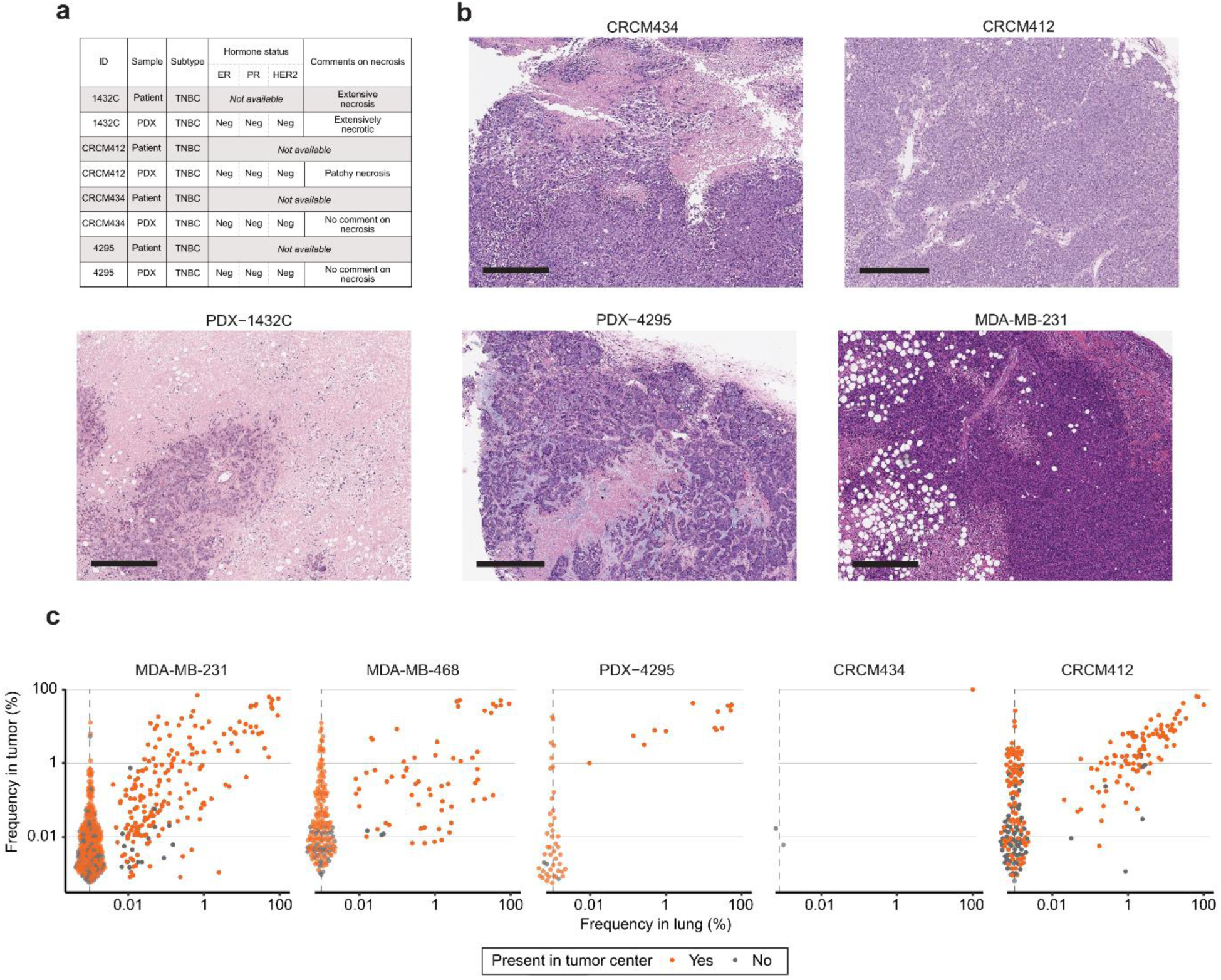
**a)** Breast cancer subtype and hormone receptors status (based on pathology report) from patient samples and matching PDXs. Estrogen receptor (ER), Progesterone receptor (PR) Human epidermal growth factor receptor 2 (HER2). Comments on necrosis were shown as per pathology report, when available for patients (pathology report only available for patient 1432) and PDXs (non-barcoded primary tumours from previous PDX passages were used for analysis). **b)** H&E staining of primary tumors, scale bar 400µm. **c)** Clonal relationship between barcodes detected in primary tumors and lungs. The colour indicates the localization of the barcodes. If the barcodes were found in the center of the primary tumor (not exclusively), the dots are orange. If the barcodes were localized exclusively in peripheral pieces, the dots are grey. Each dot represents a barcode. Dots on dashed lines represent barcode uniquely found in tumor and not detected in lung. MDA-MB-231 n=9, 3 independent experiments, MDA-MB-468 n= 5, one experiment, PDX-4295 n=3, one experiment, CRCM434 n=1, 4 one experiment, CRCM412 n= 7, 3 independent experiments.

**Supplementary Figure 3:**
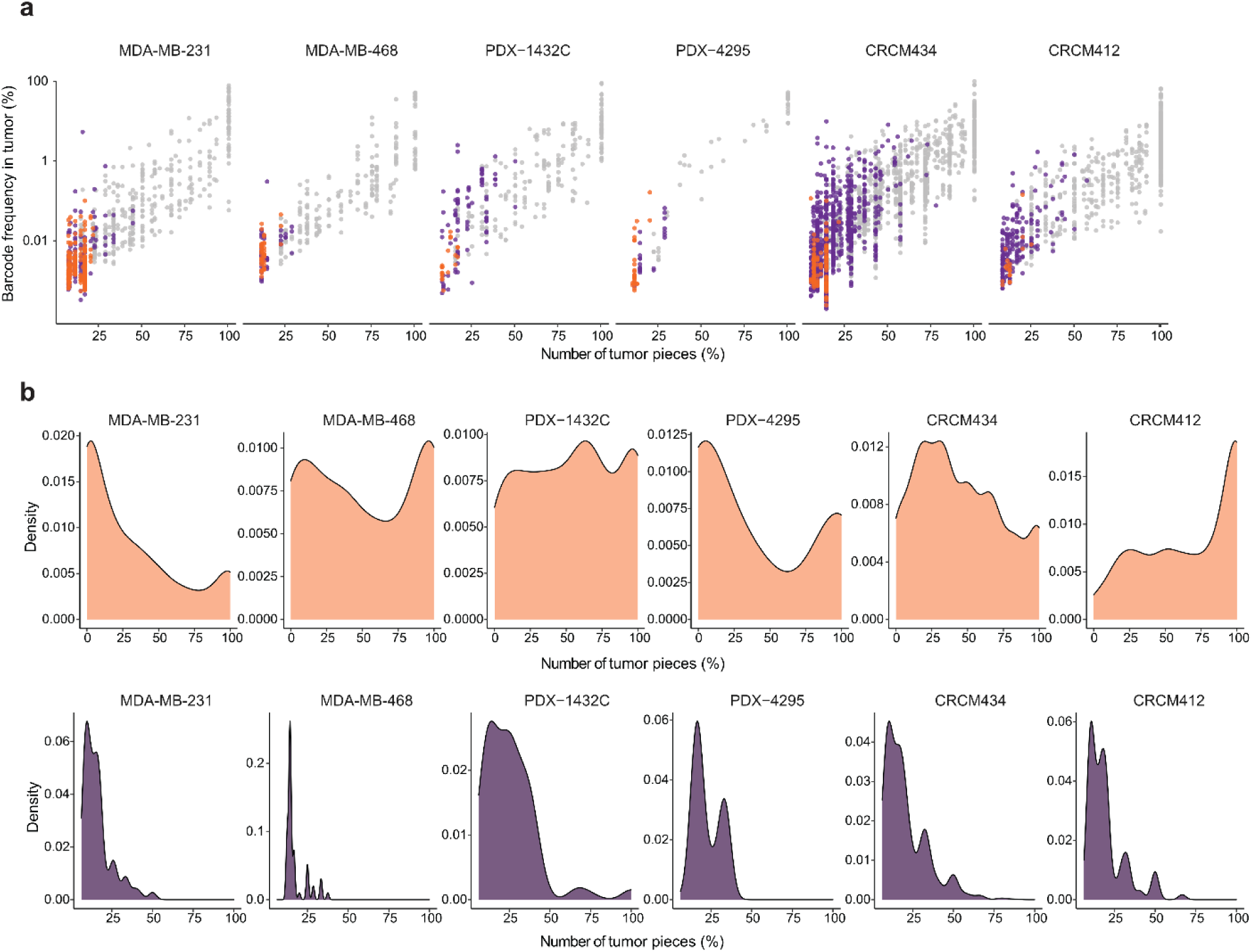
**a)** Relationship between the frequency of individual barcodes in primary tumors and the number of pieces (in percentage) containing these barcodes. Each dot represents a barcode uniquely found in the tumor center (orange), periphery (purple) or found in center and periphery (grey). **b)** Density plot showing the distribution of the number of pieces (in percentage) from the periphery containing barcodes present in the center (top), or barcodes uniquely found in peripheral pieces (bottom). MDA-MB-231 n=13, 3 independent experiments, MDA-MB-468 n= 6, one experiment, PDX-1432C n=10, 2 independent experiments, PDX-4295 n=6, one experiment, CRCM434 n=22, 4 independent experiments, CRCM412 n= 16, 4 independent experiments.

**Supplementary Figure 4:**
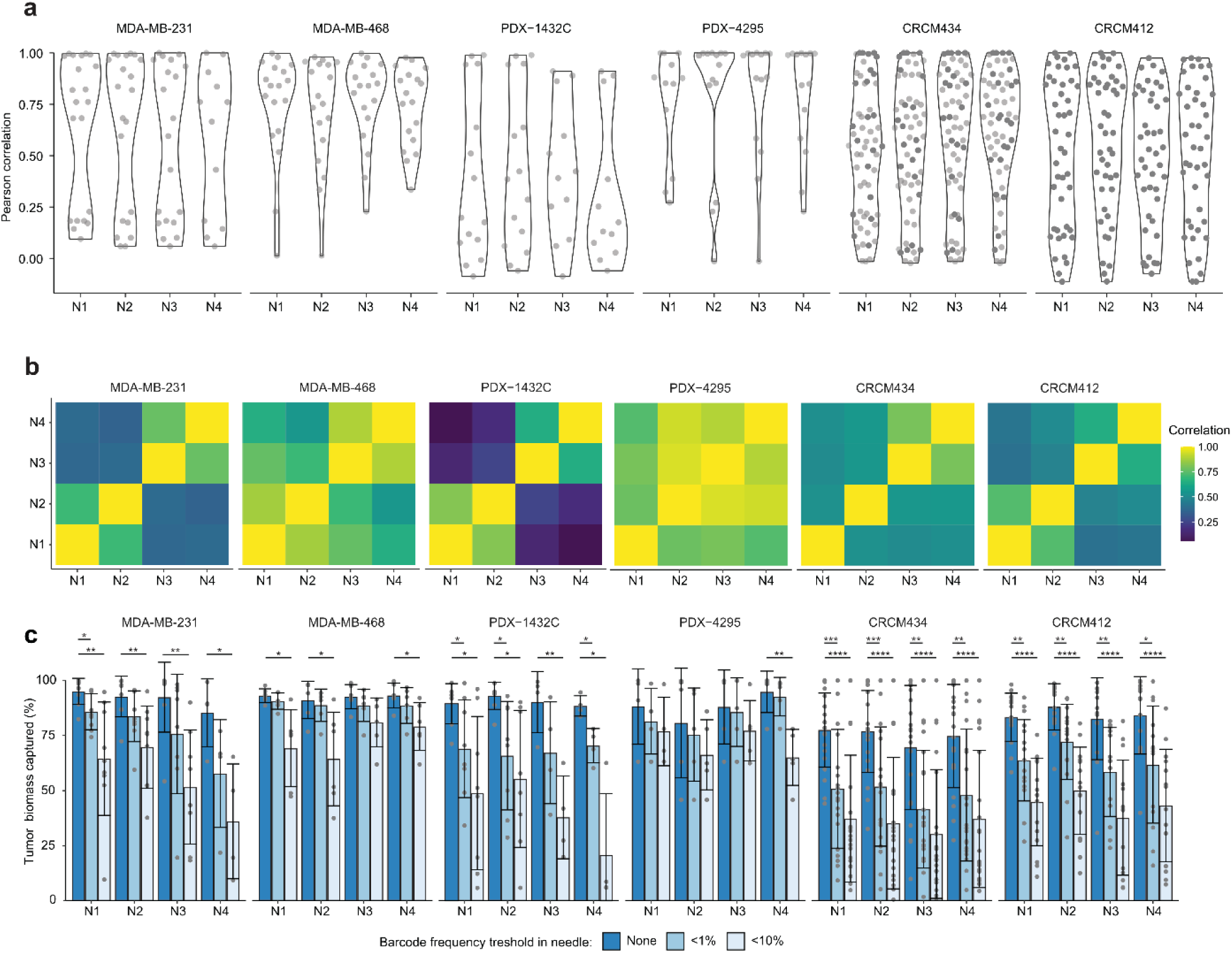
**a)** Correlation of clonal frequencies between needle samples and matching primary tumors. **b)** Pearson correlation mean comparison between different needle samples from matching tumors represented as heatmap. **c)** Percentage of primary tumor biomass captured in needle samples without threshold (left blue bar), for barcode frequency above 1% (middle bar), or above 10% (right bar). Student’s unpaired t-test, ns: non-significant= p-value>0.05, *=p-value < 0.05, **= p-value <0.005, *** = p-value <0.0005 and ****=p-value<0.0001. Error bars represent standard deviation (SD). Each dot corresponds to a needle biopsy, MDA-MB-231 n=9, 3 independent experiments, MDA-MB-468 n= 6, one experiment, PDX-1432C n=8, 2 independent experiments, PDX-4295 n=5, one experiment, CRCM434 n=22, 4 independent experiments, CRCM412 n= 16, 4 independent experiments.

**Supplementary Figure 5:**
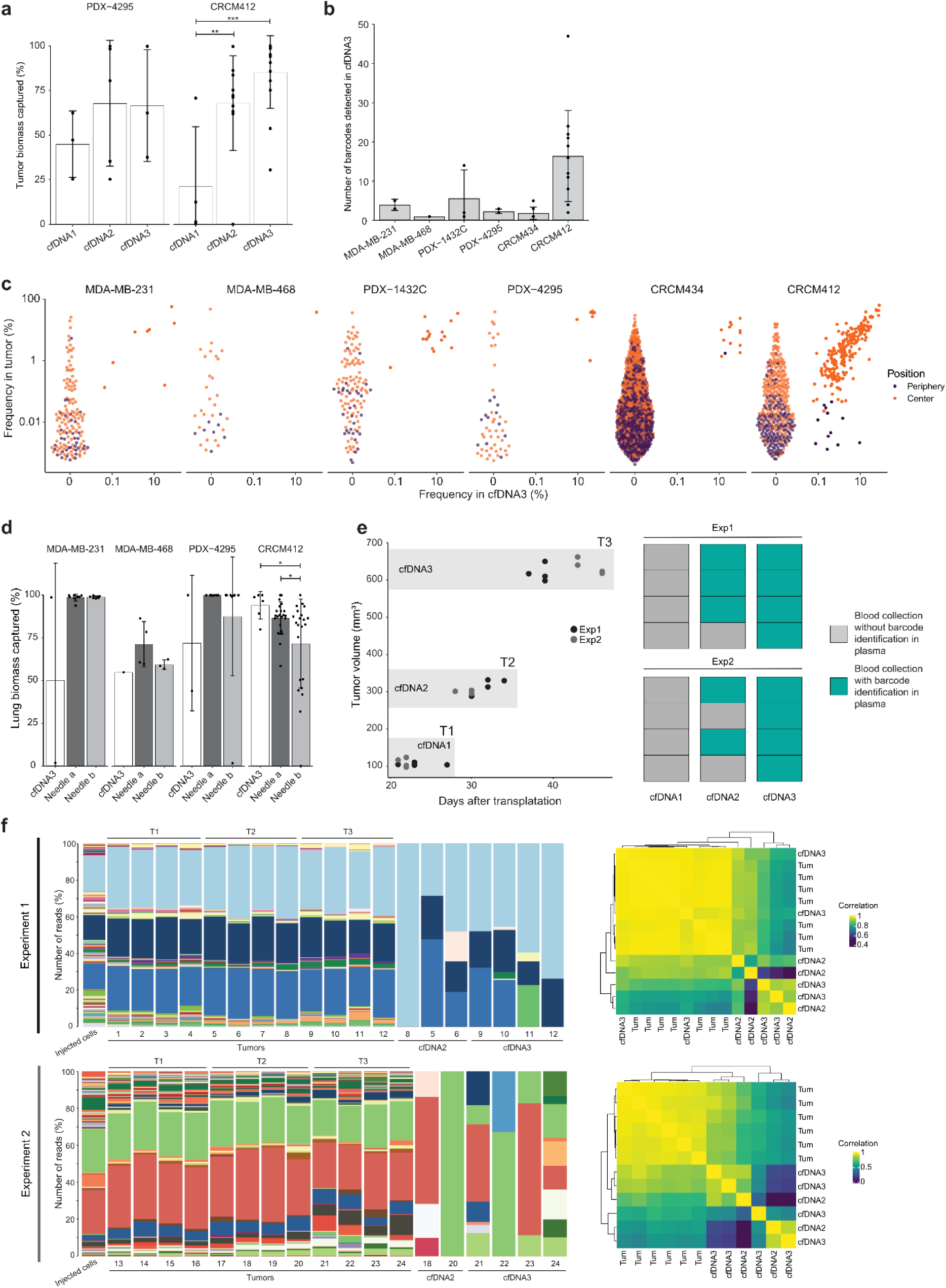
**a)** Percentage of tumor biomass captured at different timepoints. One-way Anova followed by Tukey multiple comparisons test, **= p-value <0.005, *** = p-value <0.0005. Error bars represent standard deviation (SD), PDX-4295 n=5, one experiment, CRCM412 n= 15, 4 independent experiments. **b)** Number of barcodes detected in cfDNA3 in different models. **a-b)** Each dot corresponds to a cfDNA sample. **c)** Relationship between the barccode in primary tumor and cfDNA3. Each barcode is represented by a dot, and its color represents its location in the primary tumor. Barcodes exclusively detected in peripheral pieces are represented in purple and barcodes detected in the center are represented in orange.MDA-MB-231 n=2, one experiment, MDA-MB-468 n=1, one experiment, PDX-1432C n=3, 2 independent experiments, PDX-4295 n=3, one experiment, CRCM434 n=7, 3 independent experiments, CRCM412 n= 14, 4 independent experiments. **d)** Lung cancer cell biomass captured with cfDNA3 samples, deep needle samples (Needle a) or shallow needle samples (Needle b). One-way Anova followed by Tukey multiple comparisons test, *= p-value <0.05. Error bars represent standard deviation (SD). Each dot corresponds to a biopsy sample. **e)** Tumor volume of tumors PDX-1432C at the time of harvesting for terminal end bleed (left). Each dot corresponds to a tumor. Barcode detection in plasma at the different time point, success (blue box), unsuccessful (grey box) (right panel). **f)** Representation of the clonal composition in tumors from clone splitting experiment (left panels) and their correlation between primary tumor and cfDNA (right panels).

**Supplementary Figure 6:**
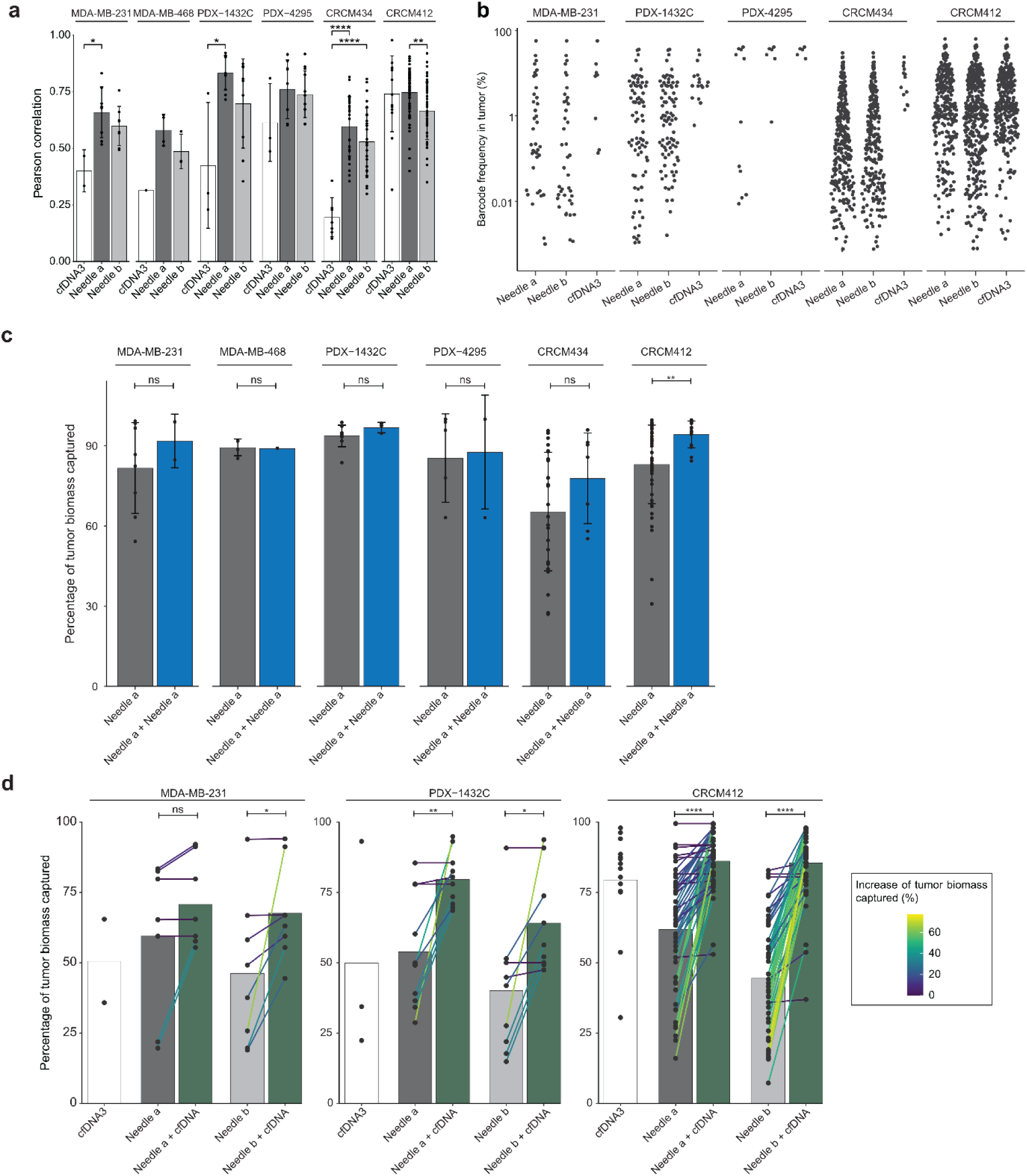
**a)** Pearson correlation coefficients measuring the barcode similarity between primary tumors and biopsies. One-way ANOVA followed by Tukey multiple comparisons test. Only significant p-value displayed, p-value > 0.05, * = p-value <0.05, **= p-value <0.005, *** = p-value <0.0005, ****= p-value <0.0001. Error bars represent standard deviation (SD). **b)** Representation of the barcode frequency in primary tumors, for barcodes detected in Needle a (deep needle samples), Needle b (shallow needle samples) and liquid biopsy (cfDNA3). Each dot represents a barcode captured in a biopsy sample. **c)** Primary tumor biomass captured in each needle sample or in two combined needle samples (blue bars). Student’s unpaired t-test, ns: non-significant, p-value > 0.05, **= p-value <0.005. Error bars represent standard deviation (SD). **d)** Primary tumor biomass captured with biopsy methods with barcode frequency in biopsy thresholds at >1%. Combination of deep needle sample and cfDNA (Na + cfDNA) or shallow needle sample and cfDNA (Nb + cfDNA). Correlated needle samples are linked with their associated increased value when cfDNA is added, and the colour of the line is scaled on the increase in tumor biomass captured as percentage. Each dot corresponds to a biopsy sample. Paired t-test, ns: non-significant, p-value > 0.05, *= p-value <0.05, **= p-value <0.005, ****= p-value <0.0001. Error bars represent standard deviation (SD). MDA-MB-231 n=1, one experiment, MDA-MB-468 n=1, one experiment, PDX-1432C n=3, two independent experiments, PDX-4295 n=3, one experiment, CRCM434 n=7, 3 independent experiments, CRCM412 n= 14, 4 independent experiments.

**Supplementary Figure 7:**
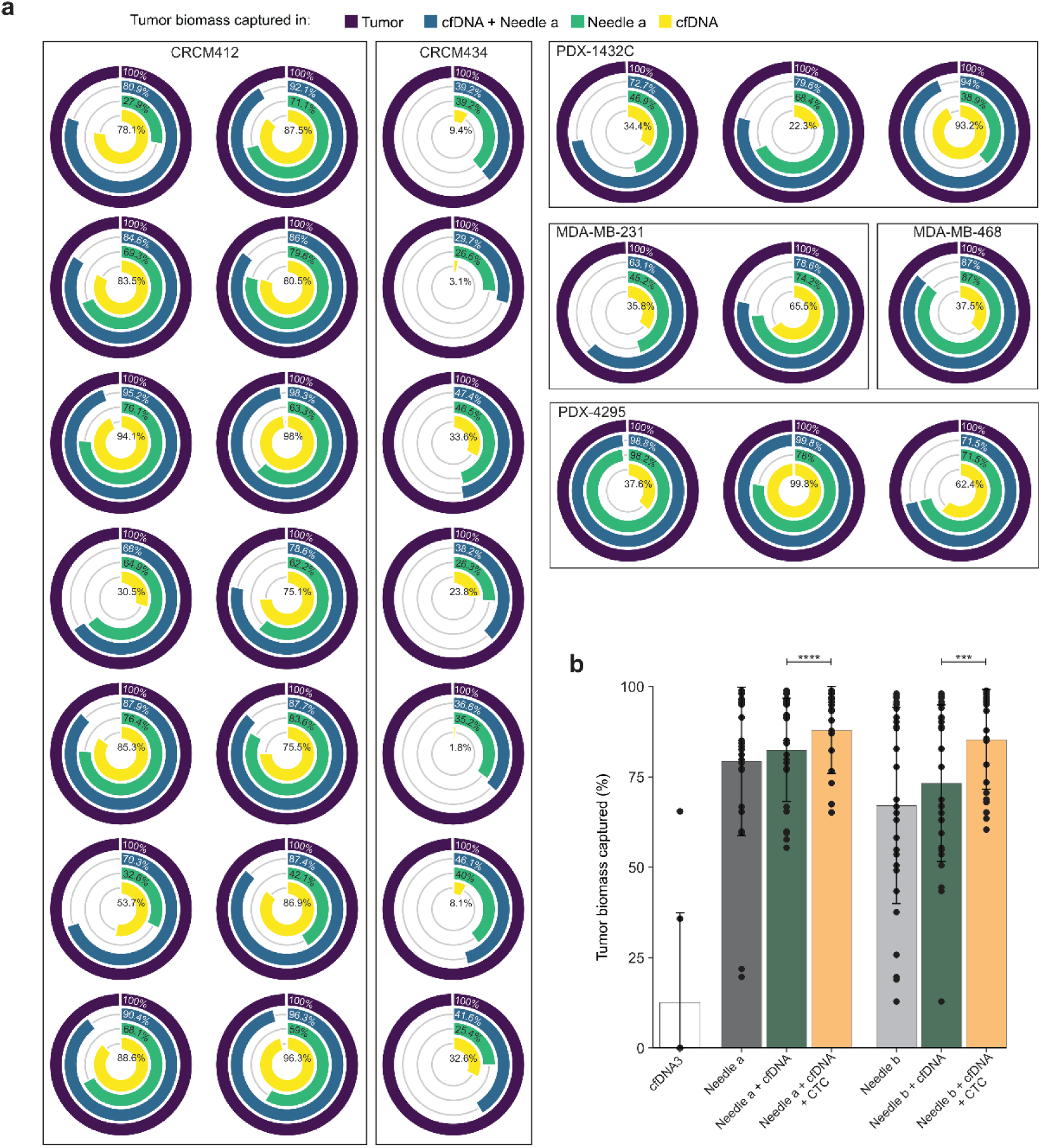
**a)** Circle plots representing the tumor biomass captured by each biopsy sampling alone or in combination, from multiple mice per models. Primary tumor plotted as reference (100%) in the outer circle (purple), inner circles represent the primary tumor biomass captured with cfDNA (yellow), deep needle samples (green) and combination of the two methods (blue). Only barcodes above 1% of the total frequency of biopsy samples were included in the computation of primary tumor biomass. **b)** Primary tumor biomass captured with biopsy methods in MDA-MB-231, with barcodes frequency in biopsy thresholds at >1%. Yellow bars indicate combinations of needles samples with cfDNA and CTCs. Paired t-test *** = p-value <0.0005, ****= p-value <0.0001. Error bars represent standard deviation (SD). MDA-MB-231 n=2, one experiment, MDA-MB-468 n=1, one experiment, PDX-1432C n=3, 2 independent experiments, PDX-4295 n=3, one experiment, CRCM434 n=7, 3 independent experiments, CRCM412 n= 14, 4 independent experiments.

